# Respiratory disease and lower pulmonary function as risk factors for subsequent dementia: a systematic review with meta-analysis

**DOI:** 10.1101/602193

**Authors:** Tom C. Russ, Mika Kivimäki, G. David Batty

## Abstract

**Background:** In addition to affecting the oxygen supply to the brain, pulmonary function is a marker of multiple insults throughout life (including smoking, illness, and socioeconomic deprivation). By meta-analysing existing studies, we tested the hypothesis that lower pulmonary function and respiratory illness are linked to an elevated risk of dementia.

**Aims:** To review the best available evidence, taken from longitudinal studies, for pulmonary function and respiratory disease as risk factors of dementia.

**Method:** We conducted a systematic review of longitudinal studies using PubMed until April 1^st^, 2019 and, where possible, pooled results in random-effects meta-analyses.

**Results:** We identified eleven studies relating pulmonary function to later dementia risk, and eleven studies of respiratory illness and dementia (including one which studied both). The lowest quartile of lung function measure Forced Expiratory Volume in one second (FEV_1_) compared with the highest was associated with a 1.5-fold (1.51, 95%CI 0.94-2.42) increased dementia risk (N_total_=127,710, 3 studies). Respiratory illness was also associated with increased dementia risk to a similar degree (1.54, 1.30-1.81, N_total_=288,641, 11 studies).

**Conclusions:** Individuals with poor pulmonary function are at increased risk of dementia. The extent to which the association between poor pulmonary function and dementia is causal remains unclear.

## INTRODUCTION

The considerable public health and care burden of dementia is well documented.(1) While the age-standardised prevalence and incidence of dementia may be declining,(2-4) because of population ageing, the absolute number of people with dementia worldwide is projected to triple from approximately 44 million in 2013 to 135 million by 2050.(5) Recent studies have provided promising findings for pulmonary function as a potential modifiable risk factor for late-life dementia. Pulmonary function affects the oxygen supply to the brain. Low pulmonary function and pulmonary disease, in turn, are associated with exposure to multiple insults across the full life course – notably smoking, illness, and socioeconomic deprivation, stunting – suggesting that they may capture a number of dementia risk factors.(6, 7) As the number of studies on pulmonary function and dementia increases, we provide the first aggregation of these results by conducting a systematic review and meta-analysis of the evidence from longitudinal studies to examine the hypothesis that low pulmonary function and pulmonary disease are risk factors for later dementia.

## METHODS

In accordance with the PRISMA guidelines,(8) we searched PubMed for articles reporting longitudinal (cohort) studies linking pulmonary function or respiratory illness with dementia occurrence from the inception of the database (1951) until 1^st^ April 2019. The search strategy combined the terms dementia OR alzheimer* AND “forced expiratory volume” OR “expiratory volume” OR FEV OR “forced vital capacity” OR “vital capacity” OR FVC OR “peak expiratory flow” OR “peak flow” OR PEF OR ((pulmonary OR lung OR respiratory) AND function) OR asthma OR COPD OR “respiratory disease” or COAD or “airways disease” OR “lung disease” OR pneumonia AND longitudinal OR prospective OR cohort. We also scrutinised the reference sections of retrieved papers and searched our own files. TCR screened the search results using Covidence (https://www.covidence.org/) and extracted data from included articles. The review protocol was registered with PROSPERO (https://www.crd.york.ac.uk/prospero/; CRD42019130376).

We included studies that were: published in English; had a prospective cohort study design with individual level exposure and outcome data, including an appropriate exposure comparator; examined the effect of pulmonary function or pulmonary disease; reported dementia as an outcome; and reported either estimates of relative risk (RR), odds ratios (OR), or hazard ratios (HR) with 95% CIs, or provided sufficient results to calculate these estimates.

We extracted the following information from each eligible article: name of the first author, start of the follow-up for dementia (year), study location (country), number of participants, number of dementia events, mean follow-up time, mean age of participants, proportion of women, method of dementia ascertainment, and covariates included in the adjusted models. The methodology of each study informed an assessment of risk of bias. Meta-analyses were conducted using R for Windows 3.4.0 and the metafor and forestplot packages. In preliminary analyses, heterogeneity measured by the I^2^ statistic was not consistently low (range: 0-94%) and so random effects models were used.

## RESULTS

Our search returned 673 articles of which 627 were discarded after review of their abstract and/or title; 46 were read in full (Figure 1). Of these, 25 studies were excluded (eight did not focus on pulmonary function or disease, six were cross-sectional, six did not have dementia as an outcome, two were ongoing studies and no results had been published, three articles were commentaries, and one study had no comparator) with the remaining 21 being included in further analyses. We considered the results for studies recording pulmonary function (N=11) and respiratory illness (N=11) separately (one study reported both pulmonary function and respiratory illness).

**Figure 1.**
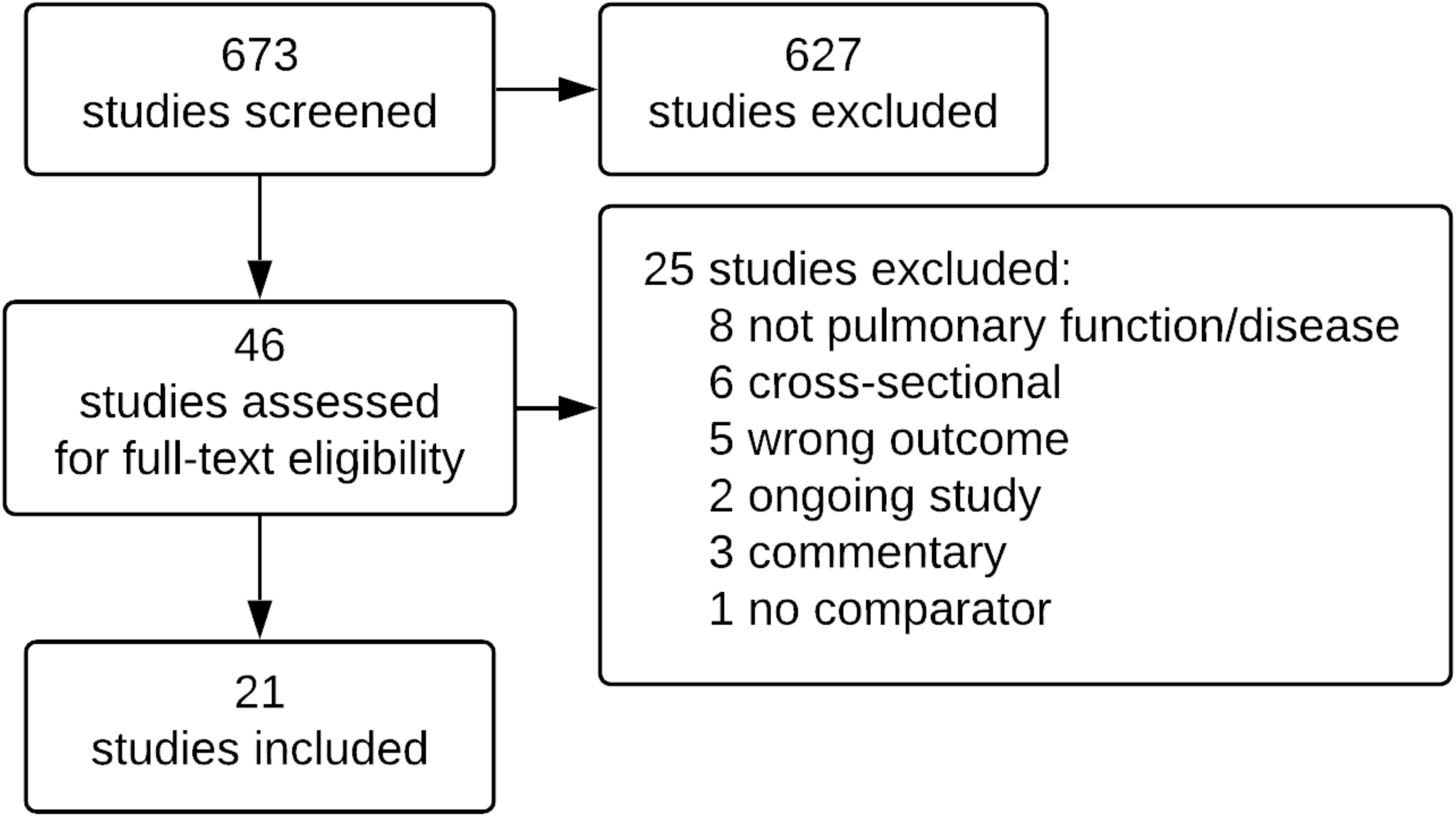
PRISMA flowchart

### Pulmonary function as a risk factor for dementia

We identified eleven prospective cohort studies that have been used to examine the association between pulmonary function and later dementia (Table 1). Mean age of participants when respiratory function was measured varied between 40 and 65 in eight studies(9-16) and was over 65 in three studies.(17-19) In these studies investigators used one of several spirometric measures as the exposure of interest (risk factor): peak expiratory flow (PEF) refers to the maximum speed of forced expiration (in litres per second); Forced Expiratory Volume (FEV) denotes the volume of air (in litres) which can be expired in a specified period of time, usually in one second (FEV_1_); and Forced Vital Capacity (FVC) captures the total volume of air which can be expired. Maximal inspiration is required before each measurement and many studies allow a defined number of attempts and record the best performance. These measures of pulmonary function correlate closely with each other(14) suggesting that associations seen for one measure with dementia/cognitive impairment are likely to be replicated in the other measures.

**Table 1.**
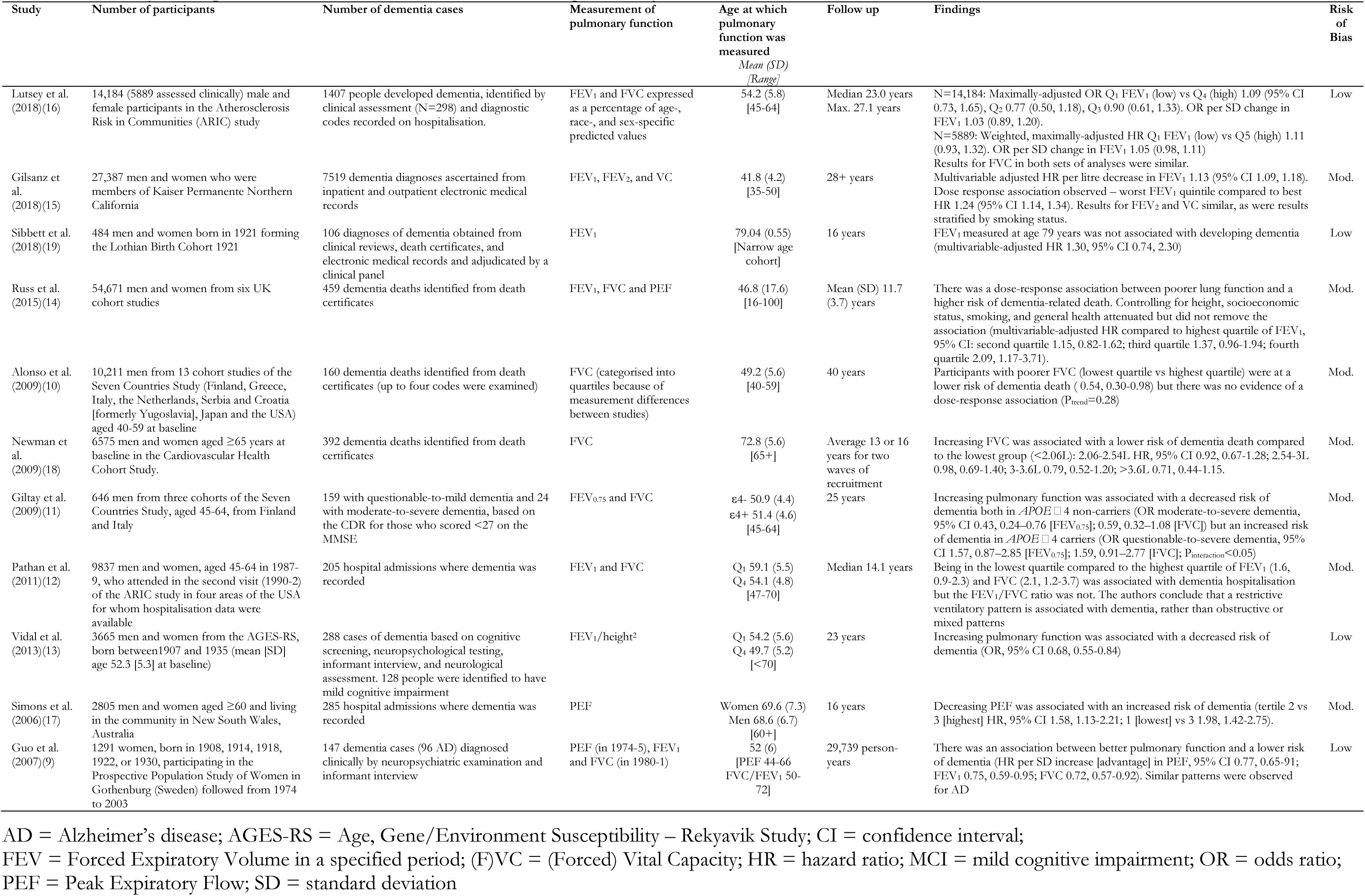
Summary of longitudinal studies of the association between pulmonary function and dementia

A range of methods were used to ascertain dementia, some in combination: death certification,(10, 14, 18, 19) linkage to electronic medical records (e.g., hospital discharge records),(12, 15-17, 19, 20) and clinical assessment.(9, 11, 13, 16, 19) A number of these studies were originally instigated to investigate risk factors for cardiovascular disease(10-12, 16, 18) or the menopause(9) and then repurposed to include dementia follow-up as study participants aged; only two were specifically set up to study diseases of ageing.(13, 19) Duration of follow up ranged from 12 to 40 years.(10, 14)

Figure 2 shows two meta-analyses of studies on FEV and dementia risk – one for categories of FEV and one for a unit change. Only three studies (127,710 adults, 1905 dementia cases) compared the lowest quartile of FEV_1_ with the highest quartile;(12, 14, 16) pooling these results in a meta-analysis gave a HR of 1.51 (95% CI 0.94, 1.64; P=0.092; I^2^=47.0%). Pooling the five studies which reported the effect of one standard deviation decrease (disadvantage) in FEV_1_ resulted in a HR of 1.28 (1.03-1.60; P=0.028; I^2^=79.3%; N=67,505, 2280 dementia cases). One of these studies standardised FEV_1_ by dividing by height^2^(13) but a sensitivity analysis excluding this transformation gave a similar pooled result (1.24, 0.92-1.68).(9, 11, 14, 16).

**Figure 2.**
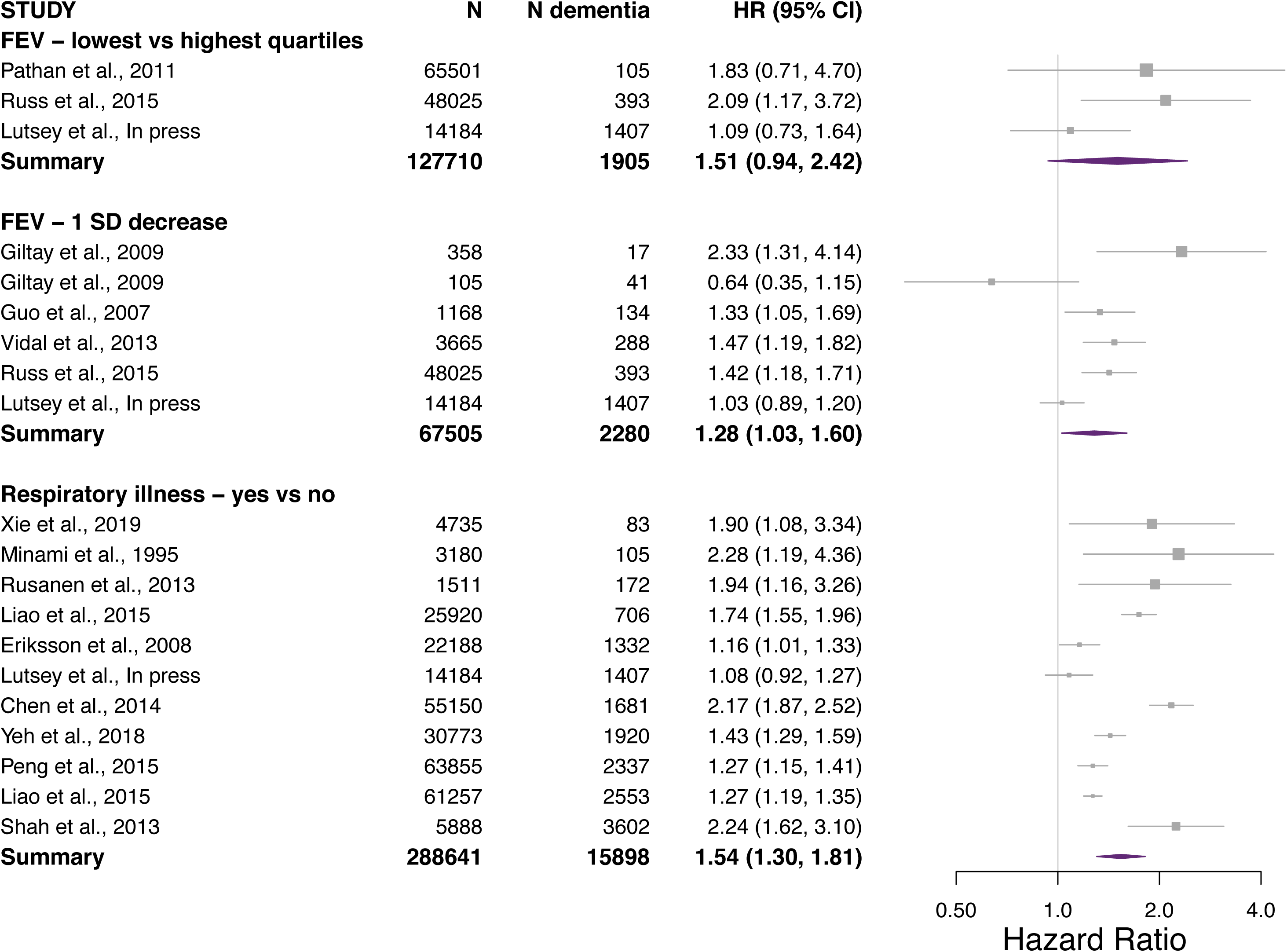
The relation of forced expiratory volume and respiratory illness with dementia — meta-analysed results

Supplementary Figure 1 shows pooled results for FVC: lowest-to-highest quartile HR 1.63 (95% CI 1.14-2.32; P=0.007; I^2^=49.3%; five studies, N=145,409, 2456 dementia cases);(10, 12, 14, 16, 18) per standard deviation disadvantage 1.21 (0.97-1.51; P=0.086; I^2^=70.7%; four studies, N=63,840, 1992 dementia cases).(9, 11, 14, 16) One study included in the latter meta-analysis reported an interaction between *APOE* status and the association between FEV/FVC and dementia which contributed to the high heterogeneity observed here; excluding this study did not affect the results of the meta-analysis but reduced the heterogeneity slightly to I^2^=65.1%.(11)

Supplementary Figure 2 shows the result of pooling two studies which compared dementia risk in the lowest vs highest quartiles of PEF, giving a HR of 2.21 (95% CI 1.73-2.82; P<0.001; I^2^=0.0%; N=50,830, 678 dementia cases);(14, 17) combining the two studies which reported the association between one standard deviation decrease in PEF and dementia gave a HR of 1.39 (1.24-1.56; P<0.001; I^2^=10.6%; N=49,316, 540 dementia cases).(9, 14)

Four studies could not be pooled in meta-analyses because of the manner in which the results were reported. A study of 27,387 Kaiser Permanente Northern California members, of whom 7519 developed dementia over more than 28 years follow up, concluded that poorer FEV_1_ (plus FEV_2_ and VC) was associated with an increased risk of dementia (multivariable-adjusted HR per litre decrease in FEV_1_ 1.13, 95% CI 1.09-1.18).(15) This finding was replicated in stratified analyses for smokers and non-smokers. Investigators in the Seven Countries Studies found that men with greater FVC were less likely to die with dementia than men with lower FVC (multivariable-adjusted hazard ratio for highest quartile [Q4] vs lowest quartile [Q1] 0.54, 95% CI 0.30-0.98) but the association observed in this study did not follow a dose-response gradient (HR, 95% CI: Q3 vs Q1 1.03, 0.63-1.68; Q2 vs Q1 0.77, 0.46-1.28).(10) An Australian study found an association between lower PEF and increased risk of subsequent hospitalisation with dementia (adjusted HR lowest vs highest tertile 1.98, 95% CI 1.42-2.75).(17) Finally, of 484 men and women from the Lothian Birth Cohort 1936, 106 adjudicated dementia diagnoses were identified from multiple sources, including face-to-face clinical assessment.(19) No robust evidence was found to suggest that FEV_1_ measured at age 79 years would be associated with developing dementia (multivariable-adjusted HR per litre/second increase 1.30, 95% CI 0.74-2.30).

### Respiratory disease as a risk factor for dementia

We identified eleven prospective studies in which investigators had explored the association between respiratory disease and later dementia (Table 2). Mean age at which disease was ascertained varied from 50.6 to 82.9 years but was over 65 years in six of the 11 studies.(21-26) Investigators identified pulmonary disease using a National Health Insurance database,(21, 22, 25, 27, 28) hospitalisation data,(24) or self-report.(16, 23, 26, 29, 30) Dementia was ascertained from the Taiwanese insurance database in five studies,(21, 22, 25, 27, 28) face-to-face assessement by a clinician in four studies,(16, 23, 24, 30), cognitive test score in one,(26) and linkage to hospital discharge and death certificate data in one.(29)

**Table 2.** Summary of longitudinal studies of the association between respiratory disease and dementia

| Study | Number of participants | Number of dementia cases | Respiratory disease | Age at which disease ascertained<br>*Mean (SD)*<br>*[Range]* | Follow up | Findings | Risk of Bias |
| --- | --- | --- | --- | --- | --- | --- | --- |
| Xie et al. (2019)(26) | 4735 participants in the Chinese Longitudinal Health Longevity Survey (CLHLS) | 83 people were newly identified as having dementia, presumably based on MMSE score | Self-reported COPD diagnosis | 82.9 (9.74) | 3 years | Maximally-adjusted HR COPD vs no COPD 1.90 (1.08–3.33). In current smokers, the same HR was 3.38 (1.09–10.5). | High |
| Lutsey et al. (2018)(16) | 14,184 (5889 assessed clinically) male and female participants in the Atherosclerosis Risk in Communities (ARIC) study | 1407 people developed dementia, identified by clinical assessment (N=298) and diagnostic codes recorded on hospitalisation. | Participants were divided into four groups based on self-reported information and spirometry: normal, respiratory symptoms with normal spirometry, restrictive impairment pattern, and COPD | 54.2 (5.8)<br>[45-64] | Median 23.0 years<br>Max. 27.1 years | N=14,184: Maximally-adjusted HR COPD vs normal 1.08 (95% CI 0.92, 1.27)<br>N=5889: Weighted, maximally-adjusted OR COPD vs normal 1.16 (95% CI 0.74, 1.82) | Low |
| Yeh et al. (2018)(25) | 30,773 men and women from the Taiwanese National Health Insurance Research Database | 1920 diagnoses of dementia | >2 clinical contacts recording asthma-COPD | Asthma-COPD<br>65.6 (11.8)<br>No illness<br>65.5 (11.9) | 10 years | Asthma-COPD was associated with an increased risk of subsequent dementia (multivariable-adjusted HR 1.43, 95% CI 1.29, 1.59) | Mod. |
| Peng et al. (2015)(28) | 12,771 people with asthma and 51,084 matched controls. | 2337 individuals identified from the Taiwan National Health Insurance database. | New diagnoses of asthma recorded on the National Health Insurance database. | Asthma<br>53.8 (17.3)<br>No illness<br>53.7 (17.4) | 11 years | Asthma was associated with an increased risk of dementia (adjusted HR, 95% CI 1.27, 1.15-1.41). | Mod. |
| Liao et al. (2015)(22) | 20,492 men and women with COPD and 40,765 matched controls. | 2553 individuals identified from the Taiwan National Health Insurance database. | New diagnoses of COPD recorded on the National Health Insurance database. | COPD<br>67.0 (12.5)<br>No illness<br>68.2 (12.4) | Mean (SD) 6.3 (3.5) years for cases, 6.9 (3.4) for controls | COPD was associated with an increased risk of dementia (adjusted HR, 95% CI 1.27, 1.20-1.36). | Mod. |
| Chen et al. (2014)(27) | 11,030 adults aged >45 years with asthma and 44,120 age- and sex-matched controls. | 1681 individuals identified from the Taiwan National Health Insurance database. | Asthma recorded on the National Health Insurance database. | 60.88 (10.39) | 8.0±3.0 years | Having asthma was associated with a doubling of risk of developing dementia (adjusted HR, 95% CI 2.17, 1.87-2.52). | Mod. |
| Liao et al. (2015)(21) | 8640 men and women ≥40 years hospitalized with COPD and 17,280 age-, sex-, and admission year-matched controls. | 706 individuals with AD or Parkinson's disease identified from the Taiwan National Health Insurance database. | COPD recorded on the National Health Insurance database. | 68.76 (10.74) | Not stated | COPD was associated with an increased risk of dementia (adjusted HR, 95% CI 1.74, 1.55-1.96). | Mod. |
| Eriksson et al. (2008)(29) | 22,188 twins in the population-based Swedish Twin Registry | 1332 twins had a record of dementia from hospital discharge and death certificate data. | History of atopy (asthma, eczema, or rhinitis) | 52.9<br>[37-71] | 22.6±7.7 years | History of atopy was associated with a modest increased risk of AD (HR, 95% CI 1.16, 0.98-1.37) or dementia (1.16, 1.01-1.33). | Mod. |
| Shah et al. (2013)(24) | 5888 participants in the Cardiovascular Health Study | 3602 identified by two-phase screening, including clinical assessment | Hospitalisation with pneumonia | 72.8 (5.6) | 10 years | Pneumonia was associated with an increased risk of later dementia (HR, 95% CI 2.24, 1.62-3.11). | Low |
| Minami et al. (1995)(23) | 3180 people without dementia in Sendai, Japan, of whom 2461 were followed up. | 105 individuals were clinically diagnosed with dementia at follow up. | Self-reported respiratory disease. | ≥65 years | 3 years | Respiratory disease was associated with a doubling of risk of dementia (adjusted OR, 95% CI 2.28, 1.19-4.36) | Mod. |
| Rusanen et al. (2013)(30) | 1511 male and female participants, aged 39.2-64.1 years at baseline, followed up at either of two points (1998 and/or 2005-8) from a random sample of 2000 (at baseline: 1972, 1977, 1982 or 1987) from four cohort studies in Eastern Finland | 172 identified by screening and, for those screening positive, clinical examination | Self-reported diagnosis of COPD or asthma | 50.6 (6.0)<br>[39.2-64.1] | Mean (SD) 25.5 (6.2) years | Pulmonary disease at baseline was associated with an increased risk of later dementia (HR, 95% CI 1.94, 1.16-3.27). Pulmonary disease in 1998 was associated with a decreased risk of dementia in 2005-8 (0.42, 0.19-0.93). | Mod. |

AD = Alzheimer's disease; CI = confidence interval; COPD = Chronic Obstructive Pulmonary Disease; HR = hazard ratio;
IQCODE = Informant Questionnaire on Cognitive Decline in the Elderly ;
RBANS = Repeatable Battery for the Assessment of Neuropsychological Status; SD = standard deviation

Figure 2 shows a meta-analysis of these 11 studies with a total of 288,641 individuals and 15,898 dementia cases giving a pooled HR of 1.54 (95% CI 1.30-1.81; P<0.001; I^2^=92.4%). Although the study-specific estimates were heterogeneous, they all favoured risk factor status. Excluding a study which investigated the association between atopic illnesses(29) (asthma, eczema, or rhinitis – only one of which is likely to have a substantial effect on pulmonary function) and dementia reduced the magnitude of the effect observed (1.28, 1.03, 1.60) as well as heterogeneity (I^2^=78.2%) but did not alter our conclusion.

## DISCUSSION

Our main finding from the 21 articles included in this quantitative review is that individuals with poorer pulmonary function (whether impaired function or overt respiratory disease), particularly in midlife, are at an increased risk of developing dementia in later life. Comparing the group who performed poorest on spirometry with the best performers was associated with a 1.5-fold increase in risk of dementia. The presence of a respiratory illness was associated with a similar magnitude of increased risk.

The effect size we found is comparable to other accepted risk factors for dementia as reported in comprehensive meta-analyses in the World Alzheimer Report 2014(31) (Figure 3) where lower educational attainment, for example, was found to be associated with an 1.8-fold increased risk of incident dementia (effect estimate combining adjusted odds ratios and HRs, 95% CI 1.83, 1.63-2.05; 31 studies). Pooling 31 studies revealed that depression was associated with around a doubling in the risk of incident dementia (unadjusted effect estimate, 1.97, 95% CI 1.67-2.02) relative to people who were free of this psychological disorder, and having diabetes in late life was associated with a 1.5-fold increase in the risk of developing dementia of any type (1.50, 1.33-1.70) and a doubling of risk of developing vascular dementia (2.39, 1.92-2.98). The recent Lancet commission on dementia prevention, intervention and care similarly reported approximately 1.5-fold increased relative risk of dementia for individuals with midlife obesity, hypertension or later life smoking (Figure 3).(32)

**Figure 3.**
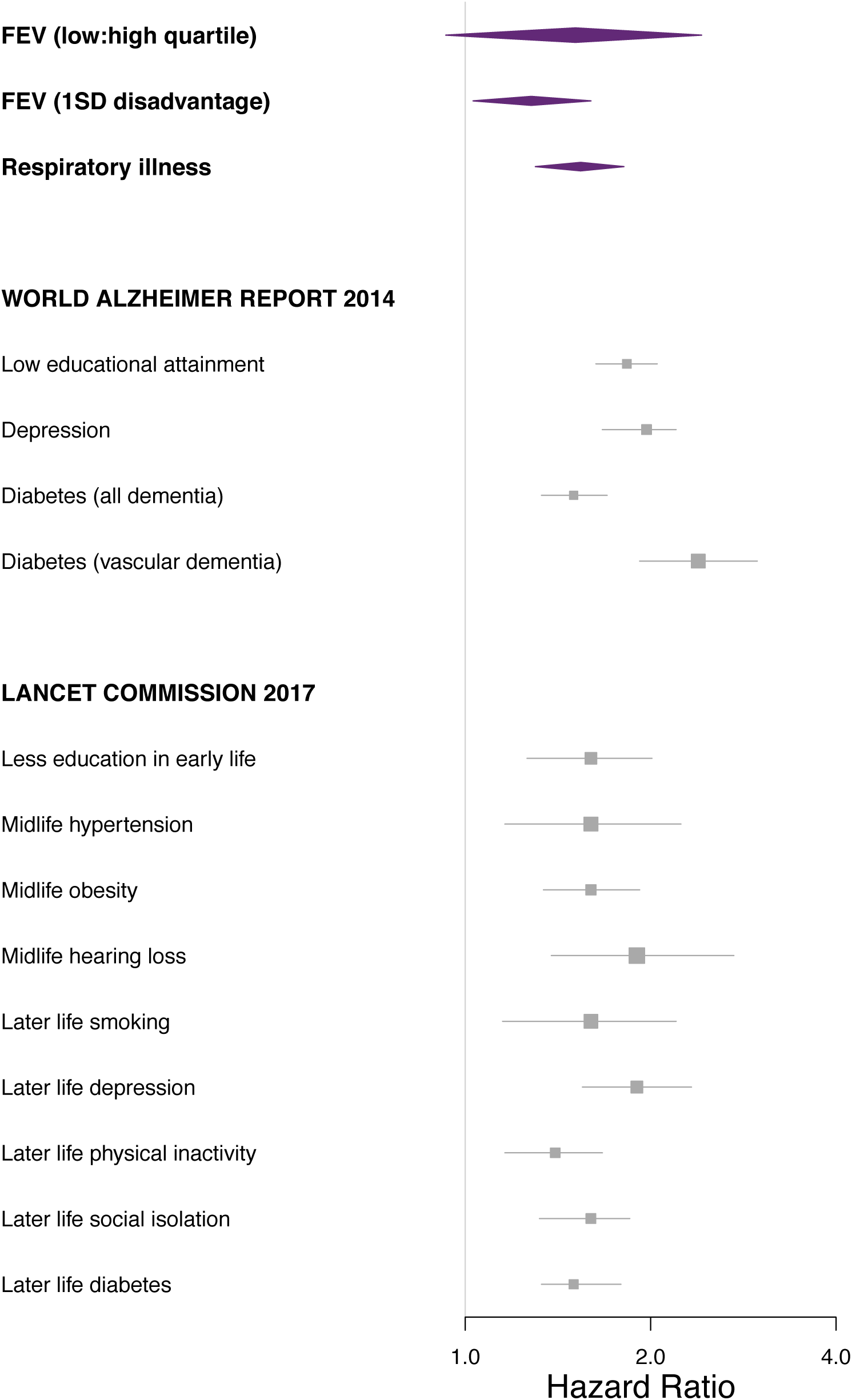
Comparison of meta-analytic findings with accepted risk factors for dementia in the World Alzheimer Report 2014(31) and Lancet Commission on dementia prevention, intervention, and care(32)

We note a recent systematic review of four longitudinal studies on the association between pulmonary function and cognitive performance.(33) While critical of the methodological quality of the studies they included, the investigators found a cross-sectional association between poorer lung function and lower levels of cognitive function, but little evidence for a longitudinal association. This may reflect different mechanisms of cognitive performance and neurological pathology underlying dementia as seems to be the case with other risk factors, such as vitamin D levels.(34, 35)

### Limitations and strengths

The comprehensive search strategy used is likely to have identified practically all relevant published studies and the inclusion of only longitudinal studies strengthens the robustness of our conclusions since, while such studies still only describe associations and do not permit causal inferences to be drawn, the temporal association between exposure and outcome adds weight to the potential importance of any identified association. The pooling of results, where possible, in random effects meta-analyses provides a quantitative summary of the evidence being a more precise estimate and overview of the literature than narrating the results of the individual studies.

There are several limitations to our work. The substantial statistical heterogeneity between studies is matched by methodological heterogeneity and – importantly – variation in the specific measure of pulmonary function used. This limited the number of studies which could be pooled in meta-analyses which may have biased our results, though the direction of bias is unlikely to be consistent in all the excluded studies.

The methodology used for dementia ascertainment is a potentially important limitation for individual studies. Face-to-face assessment by a clinician combined with brain imaging is a robust method to ascertain incident dementia cases, but is resource-intensive and differential participation in the screening process by different groups can introduce bias.(36) Linkage to electronic medical records has been shown to identify only part of a known cohort of people with dementia, particularly if multiple sources are used, although mild and undiagnosed cases are not captured.(14, 37) Death certification has been criticised as a methodology for identifying dementia cases but reporting of dementia on death certificates seems to be becoming increasingly comprehensive: for instance, a recent investigation found that, in a memory clinic cohort, of all the patients with diagnosed dementia who died during the follow-up, death certification correctly identified the diagnosis in as large a proportion as 70% of deceased patients.(38) Furthermore, investigators studying BMI and dementia found their results were similar whether dementia was ascertained solely from mortality data or whether other methods were also used.(39)

### Plausible mechanisms

A number of lines of research, in combination with the disappointing results from preventive interventions of dementia implemented at older ages,(40, 41) provide circumstantial support for aetiological process acting from across the life course.(42-46) First, recent diagnostic criteria for Alzheimer disease acknowledge a long induction period for dementia such that an asymptomatic ‘preclinical’ phase is now part of the classification.(47) Second, this accords with findings from pathological and epidemiological studies suggesting that dementia has its origins earlier in life than previously thought. For example, autopsy studies of individuals of all ages demonstrate that Alzheimer-type pathology begins to develop decades before the clinical onset of symptoms.(48-50) Third, among persons without dementia, measurements of the Alzheimer biomarker, cerebral amyloid pathology suggest a 20- to 30-year interval between first development of amyloid positivity and onset of clinial dementia.(51) Fourth, there is some evidence from prospective cohort studies that cardiovascular disease risk factors measured in midlife are associated with later dementia risk.(10, 30, 52-54) For example, in a recent analysis of the Atherosclerosis Risk in Communities (ARIC) study, midlife hypertension and elevated midlife systolic blood pressure predicted accelerated cognitive decline during 20 years of follow-up.(55) That cardiovascular disease risk factors ‘track’ from early life into adulthood(56, 57) – for example, there is a correlation between blood pressure measured in childhood and again later life(58) – is consistent with the long-term influence of exposures occurring in childhood. Fifth, a recent observational study linked *early* life cardiorespiratory fitness with later young-onset dementia,(59) though a similar association was not observed between cardiovascular disease risk factors and late-onset dementia mortality elsewhere.(60)

There are at least three plausible mechanisms for the observed association: (i) hypoxic damage to the brain resulting from poorer pulmonary function; (ii) pulmonary function may serve as a proxy for other exposures earlier in the life course which increase dementia risk; and (iii) the association may result from the shared aetiology between pulmonary, cardiovascular disease and dementia. These mechanisms will now be considered in turn.

First, the hypoxia theory proposes that poor pulmonary function is not only a risk marker but also a possible risk factor for dementia through its effects on the brain’s oxygen supply. Indeed, most dementia cases in old age are with mixed pathologies including both vascular and neuronal damage. For example, the hippocampus – an area of the brain selectively affected in Alzheimer’s disease – is particularly vulnerable to ischaemic damage,(61) although animal models of chronic hypoperfusion demonstrate impairment of spatial working memory and slowly evolving white matter abnormalities but no neuropathological changes in the hippocampus.(62) Future analyses of magnetic resonance imaging in large prospective cohort studies, such as UK Biobank(63) or the European Prevention of Alzheimer’s Disease (EPAD) Longitudinal Cohort Study(64) could help interrogate more closely the putative influence of hypoxia on the hippocampus and other areas of the brain.

Second, similarly to physical stature with which it is correlated, lung function may reflect life course exposures which modify an individual’s risk of dementia.(6, 44, 65, 66) As alluded to above, the life course paradigm in epidemiology hypothesises that exposures at different points in the life course could influence the risk of developing dementia, either through an accumulation of risk or through exposure at critical/sensitive periods.(44-46) Researchers from the Age, Gene/Environment Susceptibility study in Reykjavik, Iceland, for example, reported that smaller birth size (considered a measure of intrauterine experience) was related to poorer cognitive function at the age of 75, providing the first evidence that even the time before birth is relevant to cognitive ageing.(67) Other mechanisms includes: impaired growth leading to reduced maximal lung function; exposure to environmental factors affecting lung function and development, such as tobacco smoke (direct or indirect);(68) illness, such as childhood lower respiratory tract infections(69) and airway hyperresponsiveness;(70) socioeconomic factors (poverty, educational failure, and less-advantaged social class,(71-77); environmental factors affecting lung function, such as atmospheric pollution(72) and local exposure to traffic.(78, 79)

Third, both Alzheimer’s disease and vascular dementia may share some aetiology with cardiovascular disease and this overlap in the conditions might explain the association, independent of smoking.(80-82) It has been hypothesised that oxidative stress, inflammation, and amyloid deposition may link these two important conditions.(83, 84) In particular, oxidative stress and synaptic dysfunction appear to be closely linked(85) and brain ischaemia – which could result from cerebral atherosclerosis and stroke – causes oxidative stress-mediated damage.(86) This may possibly be exacerbated by the pro-inflammatory function of *APOE*, but this effect is controversial.(87) (88)

### Clinical and public health implications

In terms of prevention, the possibility that the association between pulmonary function and cognition might reflect a cause-and-effect relation is particularly important. To date, however, plausible mechanisms linking pulmonary function to dementia include both causal and non-causal explanations and further research on this issue is therefore needed.(9)

The point in time at which risk factors are measured seems to be important for their ability, or lack of it, to predict later dementia or cognitive decline: There was some evidence of be an age-dependent association with stronger links seen for midlife than old age respiratory function and respiratory disease. Further research is needed to clarify whether this reflects the longer exposure period among younger individuals or a critical period in which poor respiratory function is particularly damaging.

### Future directions

Pulmonary function alone is likely to have relatively low sensitivity and specificity as a predictor of cognitive decline and dementia and therefore may not be a useful predictor of dementia in the absence of a range of other predictive factors. Further research is needed to examine this. One way forward is examination of pulmonary function and pathology as a contributor to risk factor algorithms – such as the modified CAIDE risk score(54) – given the reported associations between pulmonary function and dementia which remained after adjustment for cardiovascular risk factors. To date, there is some evidence to suggest that such risk scores predict cognitive function(89) and decline,(90) although there is less evidence for prediction of dementia.(91) Therefore there is, as yet, no risk score including lung function which can be used in clinical practice.

In addition to risk stratification and early identification of risk groups, further work is also required to confirm or refute the importance of pulmonary function as a risk factor amenable to modification and thus a target for prevention. Extended follow up of studies where the initial focus was treating respiratory illness might be a pragmatic place to start. It would be difficult to conduct a powered randomised, controlled trial on this topic given the long follow-up from mid-life into later life needed and the large sample size required to obtain a sufficient number of incident dementia cases. Therefore, in order to reduce confounding and reverse causation bias, Mendelian randomization studies with genetic variants related to lung function as an instrument would provide one avenue to pursue.

Further mechanistic research is also warranted in order to test in depth plausible pathways linking pulmonary function and dementia, such as the hypoxia and vascular damage hypotheses. The global public health importance of dementia is such that researchers should pursue this promising line of research on pulmonary function.

## Contributions

TCR drafted the article and all authors revised it for critical content.

## FUNDING & ACKNOWLEDGEMENTS

TCR was supported by the Alzheimer Scotland Marjorie MacBeath fellowship from 2015-16 and is now employed in the UK National Health Service. TCR and GDB are members of both the Alzheimer Scotland Dementia Research Centre funded by Alzheimer Scotland and the University of Edinburgh Centre for Cognitive Ageing and Cognitive Epidemiology, part of the cross council Lifelong Health and Wellbeing Initiative. MK is supported by the Medical Research Council, UK (S011676, R024227), NordForsk, the Nordic Research Programme on Health and Welfare (grant #75021), the Academy of Finland (311492), and the Helsinki Institute of Life Science. All researchers are independent of funders.

The authors have no conflicts of interest.

These results were presented in preliminary form as a poster at the AAIC 2017.

---

**Supplementary Figure 1.**
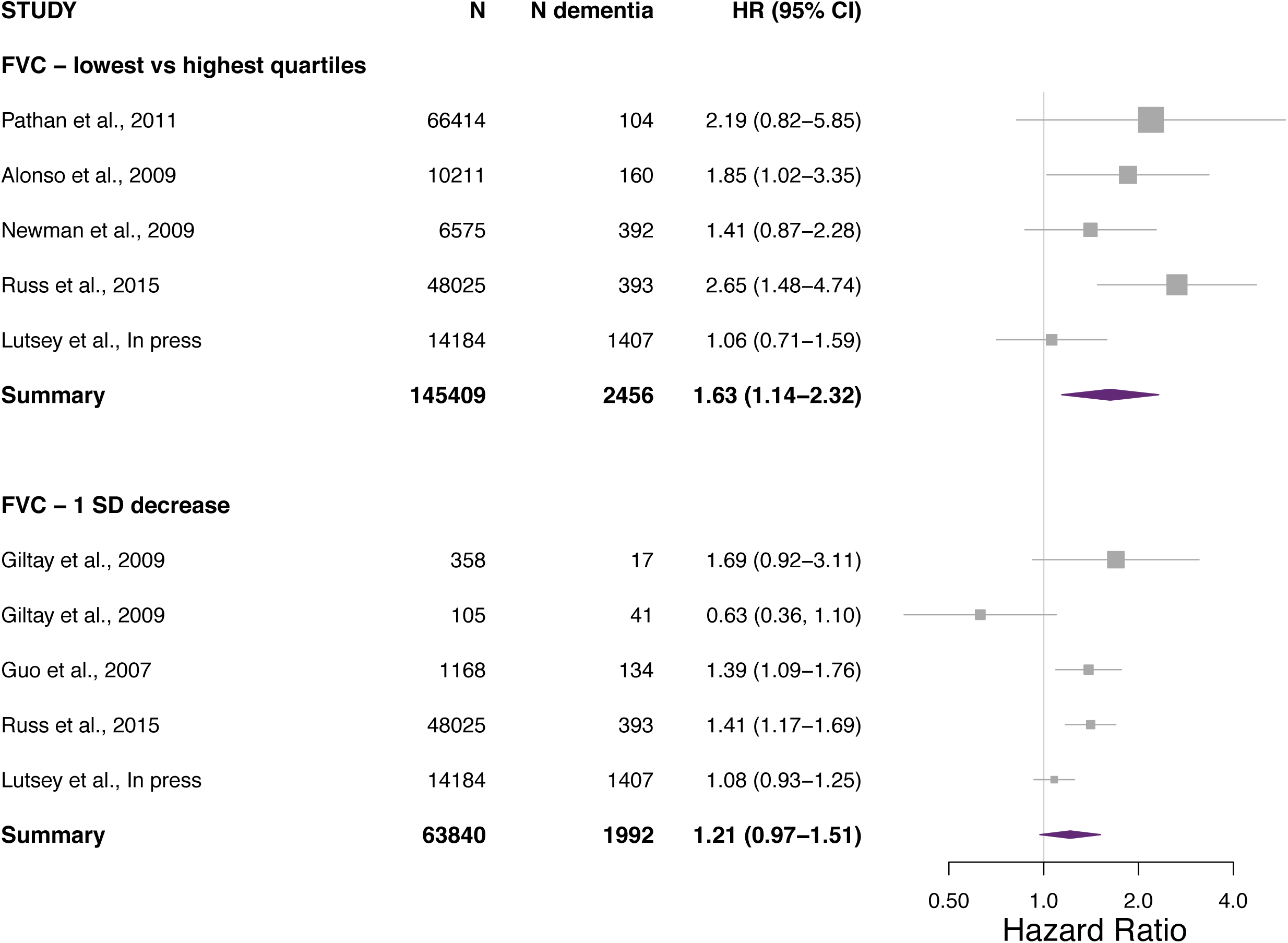
The association between Forced Vital Capacity – lowest quartile compared to highest quartile and one standard deviation decrease – and dementia with meta-analysed results

**Supplementary Figure 2.**
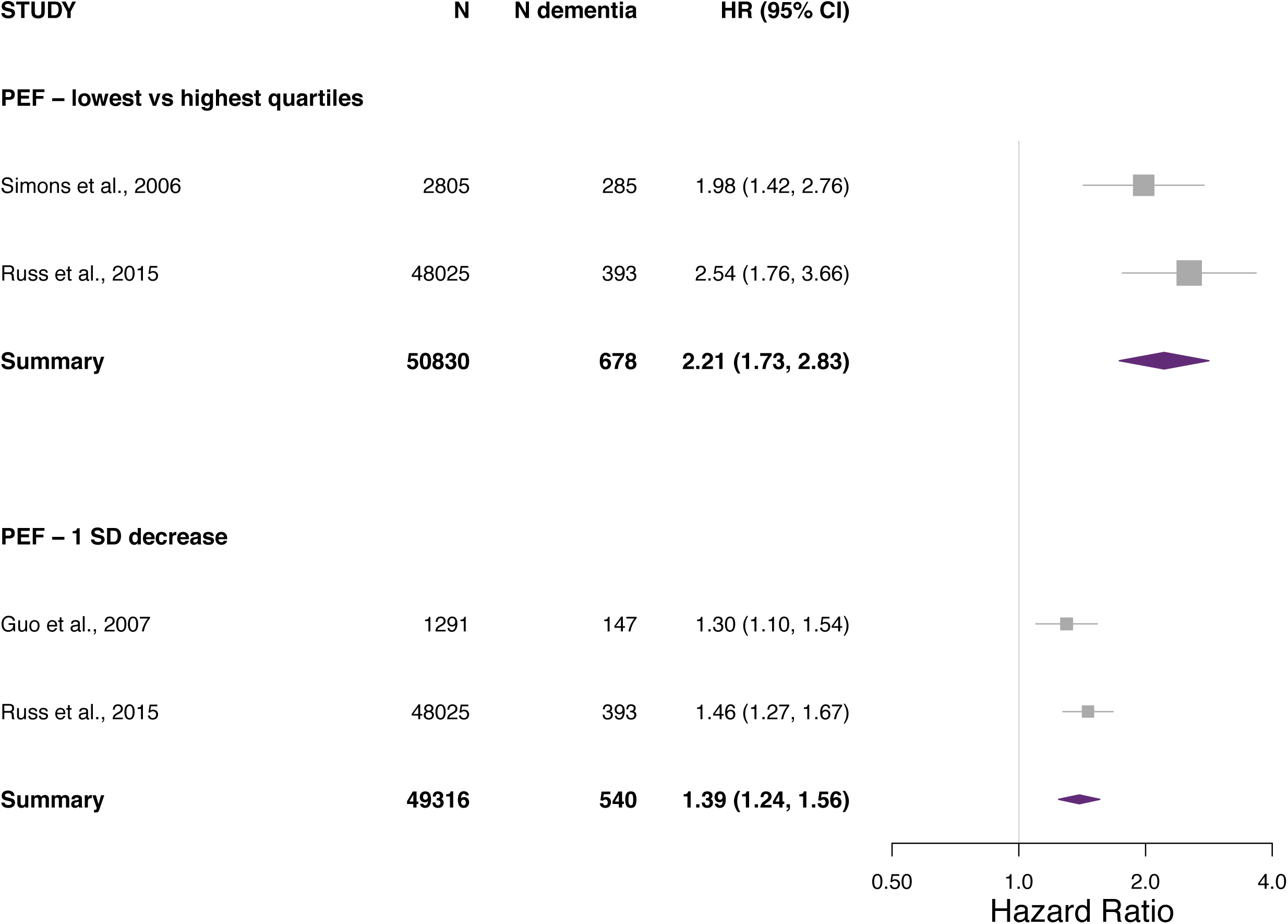
The association between Peak Expiratory Flow – lowest quartile compared to highest quartile and one standard deviation decrease – and dementia with meta-analysed results

